# Polyclonal pathogen populations accelerate the evolution of antibiotic resistance in patients

**DOI:** 10.1101/2021.12.10.472119

**Authors:** Julio Diaz Caballero, Rachel M. Wheatley, Natalia Kapel, Carla López-Causapé, Thomas Van der Schalk, Angus Quinn, Claudia Recanatini, Basil Britto Xavier, Leen Timbermont, Jan Kluytmans, Alexey Ruzin, Mark Esser, Surbhi Malhotra-Kumar, Antonio Oliver, R. Craig MacLean, WP3A working group

## Abstract

Antibiotic resistance poses a global health threat, but the within-host drivers of resistance remain poorly understood. Pathogen populations are often assumed to be clonal within hosts, and resistance is thought to emerge due to selection for *de novo* variants. Here we show that pulmonary populations of the opportunistic pathogen *P. aeruginosa* are often polyclonal. Crucially, resistance evolves rapidly in patients colonized by polyclonal populations through selection for pre-existing resistant strains. In contrast, resistance evolves sporadically in patients colonized by monoclonal populations due to selection for novel resistance mutations. However, strong trade-offs between resistance and fitness occur in polyclonal populations that can drive the loss of resistant strains. In summary, we show that the within-host diversity of pathogen populations plays a key role in shaping the emergence of resistance in response to treatment.

**One sentence summary:** Antibiotic resistance evolves quickly in patients colonized by polyclonal pathogen populations.

## Main text

Antibiotic resistance in pathogenic bacteria poses a fundamental threat to human health. It is well established that antibiotic use is associated with the emergence of resistance (*1, 2*). However, the within-host drivers of resistance remain poorly understood, making it difficult to predict the emergence of resistance at the scale of individual patients (*3, 4*). This is an important problem to address, as resistant infections are associated with worse outcomes for patients (*5, 6*).

The dominant model for the within-host emergence of resistance is that resistance evolves as a result of selection for novel alleles that are acquired by *in situ* by mutation or horizontal gene transfer (*4, 7-11*). An implicit assumption of this model is that hosts are colonized by clonal pathogen populations that lack genetic variation due to due to bottlenecks that occur during transmission (*7, 12-15*). However, hosts can also be colonized by multiple strains of the same pathogen species, giving rise to polyclonal pathogen populations (*16-19*). Polyclonal populations contain both novel genetic variation that is acquired *in situ* and pre-existing variation that reflects differences between the co-colonizing strains. A key concept from evolutionary biology is that this additional source of standing genetic variation in polyclonal pathogen populations should accelerate the evolutionary response to antibiotic treatment by increasing the genetic diversity that selection acts on (*20-22*). This simple logic predicts that resistance will evolve rapidly in hosts colonized by diverse pathogen populations. In this paper we directly test this prediction using populations of the opportunistic pathogen *P. aeruginosa* sampled from critically ill patients who were enrolled in ASPIRE-ICU, an observational trial of *P. aeruginosa* infection in European hospitals (*23*).

Clinical microbiology projects usually focus on the analysis a single bacterial isolate per patient sample, making it difficult to assess the prevalence and importance of within-patient pathogen diversity. To overcome this limitation, we sequenced the genomes of 441 isolates that were collected from lower respiratory tract samples of 35 patients from 12 hospitals (Figure 1A) (Supplementary Table 1). In line with previous work (*24*), we found that *P. aeruginosa* has a non-epidemic clonal population structure, consisting of clearly differentiated Sequence Types (STs) that are separated by long branches (Figure 1B). The most prevalent ST (ST235) segregated into three sub-lineages, which diversified prior to our sampling (Figure 1C). Given this phylogeny, we considered STs and sub-lineages of ST235 to represent unique clones. Clonal diversity in patients was bi-modally distributed (Figure 1A). The majority of patients (n=23/35) were colonized by a single clone (ie “monoclonal populations”). However, approximately 1/3 of patients (n=12/35) were colonized by multiple clones (ie “polyclonal populations”), including 10/17 patients from a single hospital with a high *Pseudomonas* colonization rate (Figure 1A). Clonal diversity in these patients tended to be high (mean Simpson’s index=.37, st.dev=.15), reflecting an even distribution of clones of polyclonal populations.

**Figure 1.**
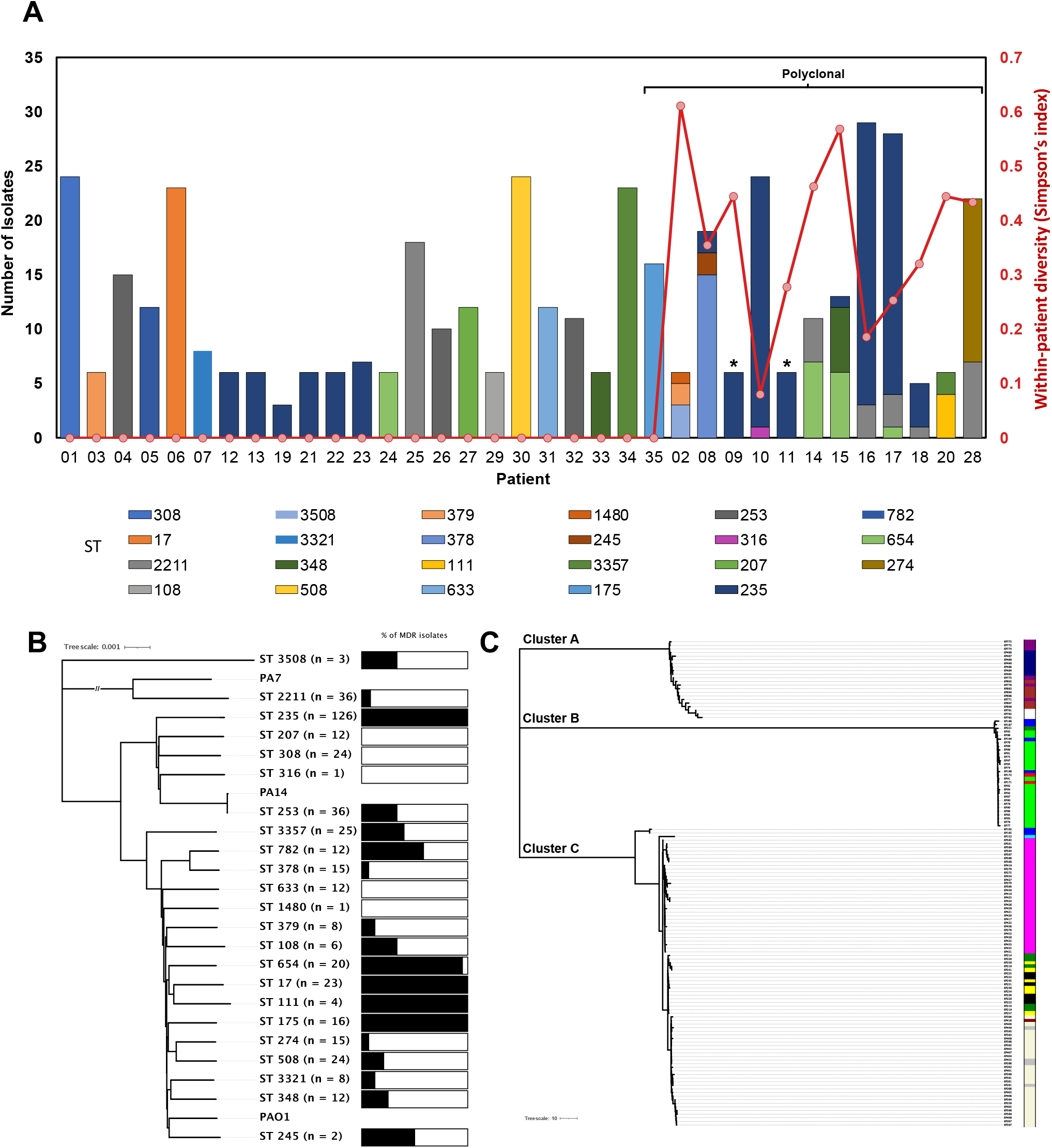
Overview of patient cohort and isolate dataset. **(A)** Within-host diversity of *P. aeruginosa*. Bars show the number of isolates and Sequence Type(s) (ST) of *P. aeruginosa* collected from each patient in study cohort. Polyclonal populations were identified in 12/35 patients. * indicates patients with a polyclonal population consisting of isolates from multiple distinct ST235 sub-lineages. Plotted points show the diversity of clones withing patients, as measured by Simpson’s index. **(B)** Neighbour joining phylogeny of all STs found in this study and the respective proportion of MDR isolates. **(C)** Neighbour joining phylogeny of ST235 isolates showing the three distinct ST235 sub-lineages (cluster A, B and C) and the patients (indicated by colour blocks) from whom they were collected. ST; Sequence Type

To test the hypothesis that within-host diversity accelerates the evolution of resistance, we measured changes in resistance over time in a sub-set of 12 longitudinally sampled patients who were treated with antibiotics that are active against *P. aeruginosa*. We measured resistance to a panel of antibiotics that included representatives of all of the major families of antibiotics (Supplementary Table 1), but was biased towards β-lactam antibiotics due to their clinical relevance for treating *Pseudomonas* infections (*25*). Changes in resistance were measured by calculating the change in proportion of isolates that were phenotypically resistant to each antibiotic over time. This approach allowed us to distinguish between direct responses to antibiotics that were used in treatment and collateral changes in resistance to antibiotics that were not used for treatment for each patient (Supplementary Table 2). Surprisingly, the average magnitude of direct and collateral responses to antibiotic treatment did not differ from each other, implying an overall tendency towards cross-resistance (F_1,67_=0.59, P=0.44). Given this, our analysis of changes in resistance in response to treatment included all antibiotics in our panel.

Antibiotics should impose strong selection for resistance in populations where average levels of resistance are low. Consistent with this hypothesis, increases in antibiotic resistance were negatively correlated with the prevalence of resistant isolates prior to treatment (Figure 2A; main effect initial resistance, F_1,67_=22.00, P<0.0001). Importantly, this effect of initial resistance did not differ between patients with monoclonal and polyclonal populations (initial resistance*diversity interaction F_1,67_=.0009, P=0.97). Response to antibiotic treatment varied between patients, even after correcting for the impact of initial resistance, implying that individual combinations of host/pathogen/treatment played an important role in the evolution of resistance (main effect of patient F_11,67_=3.90, P<0.0002). Crucially, increases in resistance were greater in patients colonized by polyclonal populations than monoclonal populations for any given level of initial resistance (Figure 2C; main effect diversity, F_1,67_=14.4, P<0.001).

**Figure 2:**
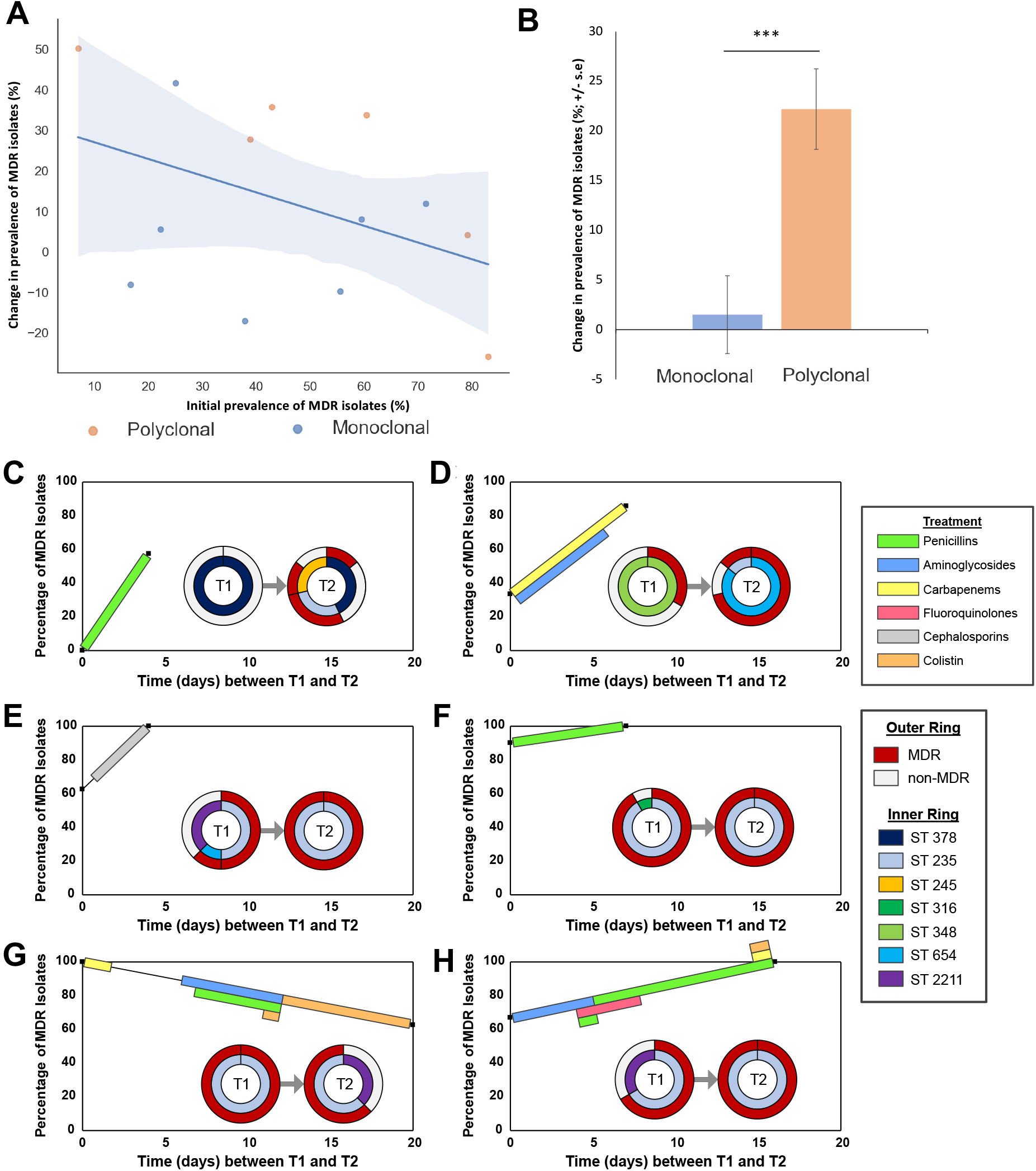
Polyclonal populations accelerate the emergence of resistance. **(A)** Change in the prevalence of MDR isolates in patients with monoclonal (blue) and polyclonal (orange) populations. Resistance emerges most rapidly when initial prevalence of MDR isolates is low (solid line). **(B)** Polyclonal populations were associated with large increases in resistance. Bars show the mean (+/- s.e) change in resistance after correcting for the effect of initial resistance (P<0.001). **(C-H)** Changes in the prevalence of MDR isolates and strain composition in longitudinally sampled patients with polyclonal populations. Percentage of MDR isolates is shown between between sampling point 1 (T1) and sampling point 2 (T2), the colour blocks between these sampling points indicate patient antibiotic use. The inset pie charts show the proportion of STs at each timepoint (inner ring), and the contribution of these STs to MDR (outer ring). (C) Patient 8, (D) Patient 15, (E) Patient 17, (F) Patient 10, (G) Patient 16, (H) Patient 18.

The accelerated evolution of resistance in polyclonal populations could have been driven by either selection for (i) novel resistance polymorphisms or (ii) pre-existing resistant strains. Examining changes in the composition of polyclonal populations revealed that non-multidrug resistant (non-MDR) STs were repeatedly replaced by ST235 and ST654, both of which are well-characterized and epidemiologically successful multidrug resistant (MDR) strains of *P. aeruginosa* (*26*) (Figure 2 C-H). Selection for these pre-existing resistant strains accounted for >90% of the increase in the prevalence of MDR isolates in polyclonal populations (Figure 3A). To complement this statistical approach, we searched for polymorphisms in known resistance genes that reflect *de novo* mutation with hosts. Resistance polymorphisms were more common in patients colonized by monoclonal populations than polyclonal populations (Figure 3B, F_1,67_=7.549, P<0.05). In contrast, genome-wide levels of polymorphisms did not differ between monoclonal and polyclonal populations, suggesting that the *de novo* evolution of resistance in polyclonal patients was not constrained by an underlying lack of genetic diversity (Supplementary Figure 1). In summary, our results show that a fundamental dichotomy exists in the mechanism of resistance evolution in patients colonized by monoclonal pathogen populations (selection for novel variants) and polyclonal populations (selection for pre-existing strains).

**Figure 3:**
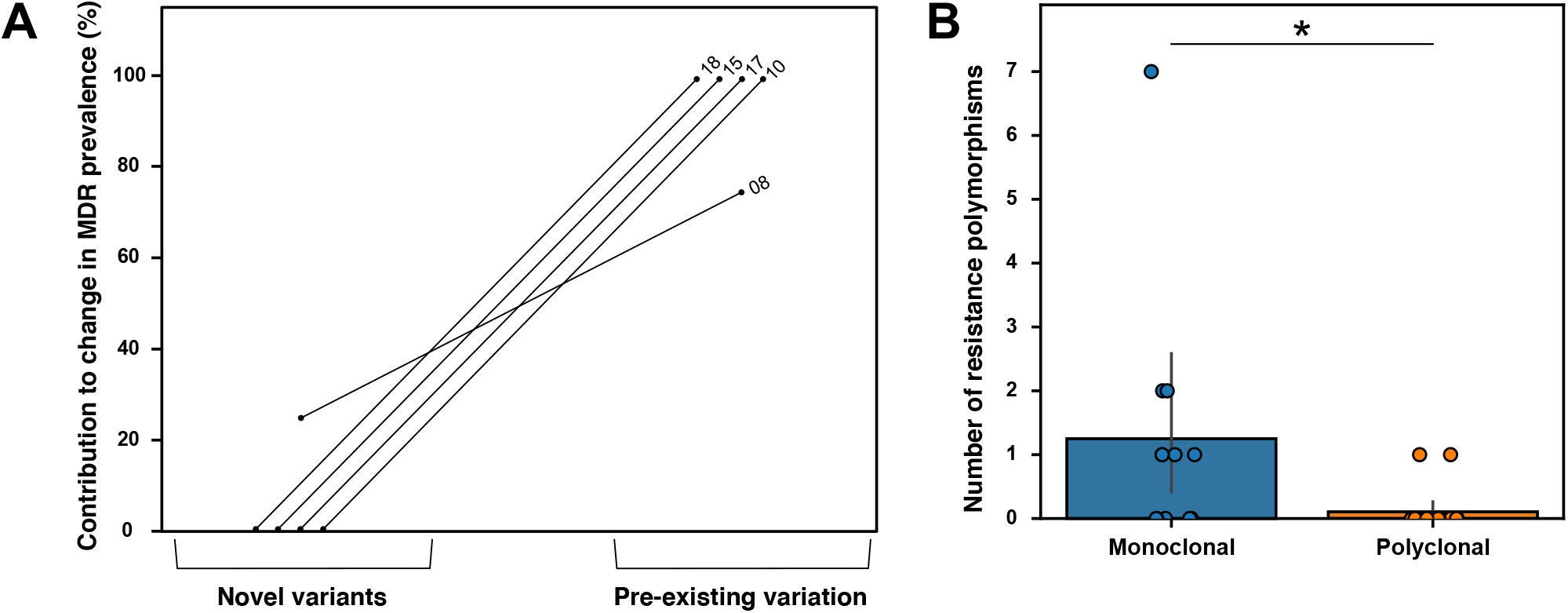
Genomic drivers of resistance evolution within patients. (**A**) Pre-existing genetic variation drives rapid evolution in polyclonal populations. We partitioned the increases in the prevalence of MDR isolates in polyclonal populations (patient numbers indicated) into changes within STs, that reflect *de novo* evolution and changes in ST composition, that reflect selection on pre-existing genetic variation. (**B**) Novel mutations drive resistance evolution in monoclonal populations. Polymorphisms in established antibiotic resistance genes that reflect *de novo* mutation were more common in monoclonal populations (P<0.05), but overall levels of genetic diversity did not differ between monoclonal and polyclonal populations (Supplementary Figure 1).

Fitness trade-offs are thought to play an important role in limiting the evolution of antibiotic resistance (*27, 28*). To test the impact of pathogen diversity on trade-offs, we measured bacterial growth rate in populations containing a mixture of MDR and non-MDR isolates (Figure 4A). MDR was associated with reduced growth, demonstrating a fitness cost to resistance (main effect MDR: F_1,167_=5.42, P=.021). Crucially, trade-offs were stronger in polyclonal populations than in monoclonal populations (MDR*diversity interaction: F_1,167_=11.24; P=0.0010), which may reflect the fact that successful MDR/XDR strains of *P. aeruginosa* typically carry a suite of chromosomal resistance mutations and horizontally acquired resistance genes (*26, 29*). Interestingly, our data set contains an example of a patient with a polyclonal population where resistance declined under antibiotic treatment due to replacement of the high resistance/low growth rate ST235 by high growth late/low resistance ST2211, highlighting the potential clinical significance of fitness trade-offs (Figure 4B). Thus, from the perspective of AMR, pathogen diversity is a double-edged sword: high diversity accelerates emergence of resistance under treatment, but accelerates the loss of resistance when antibiotic pressure is weak.

**Figure 4:**
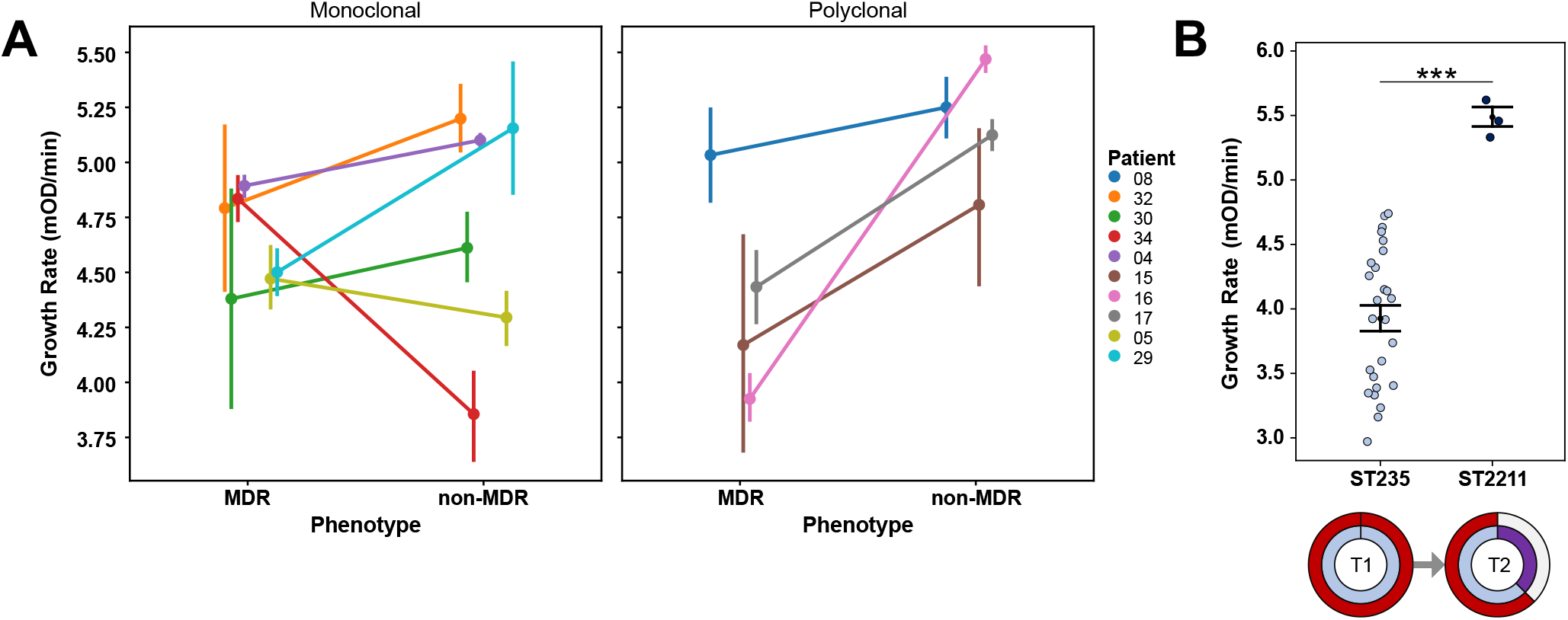
Fitness trade-offs within patients. (**A**) Polyclonal populations are associated with strong fitness trade-offs. Points show the mean growth rate in antibiotic-free culture medium (+/- s.e.m) of 179 MDR and non-MDR isolates from 10 patients. MDR was associated with reduced growth rate (P=.021), and the trade-off associated with MDR was strongest in polyclonal populations (P=.001). (**B**) Fitness trade-offs drive the loss of resistance under antibiotic treatment. The prevalence of MDR isolates declined over time in patient 16, in spite of strong antibiotic treatment, as shown in Figure 2F. Points show the comparison of the growth rate of isolates of ST235 (MDR) and ST2211 (non-MDR) collected from this patient (t_27_=4.88, P<0.0001), and the circular plot shows changes in strain composition (inner circle) and MDR phenotype (outer circle) from the first sampling point of this patient (T1) to the second (T2).

Evolutionary approaches are increasingly being used to understand and combat antibiotic resistance(*27, 30-32*), and an important challenge for this field is to understand the within-host drivers of resistance (*3, 4*). The key finding of this study is that resistance evolves rapidly in patients colonized by diverse *P. aeruginosa* populations due to selection for pre-existing resistant strains, demonstrating a clear link between within-host diversity and resistance evolution. Conventional methods used in clinical microbiology labs are systematically biased against the detection of pathogen diversity, making it difficult to assess the importance of pre-existing diversity in resistance across bacterial pathogens. For example, high levels of within-host diversity may explain why some pathogens, such as *Pseudomonas*, rapidly adapt to antibiotic treatment in patients (*33*).

Fitness costs are thought to play a key role in preventing the spread of resistance(*27*). In this case, trade-offs between resistance and growth rate make it is challenging to understand how strains that vary in resistance can stably coexist within the same patient. Therefore, we speculate that polyclonal populations will be most common in high infection rate settings, where pathogen strain diversity is high due to recurrent colonization (*10*), or in patients where antibiotic exposure is heterogeneous, allowing high and low resistance strains to effectively occupy different niches (*34*). A further challenge will be to determine if polyclonal populations arise as a consequence of single colonization events or by superinfection(*35*). In a broader context, our study underscores the importance of understanding of the drivers of within-host bacterial diversity and its consequences for within-host evolution and pathogenesis (*4, 7, 9, 36, 37*). Ultimately this may lead to novel strategies for predicting and preventing the emergence of resistance that are based on quantifying and manipulating the within host diversity of bacterial pathogens.

## Materials and Methods

### Clinical data

The subjects were recruited as part of the observational, prospective, multicentre European epidemiological cohort study ASPIRE-ICU (The Advanced understanding of *Staphylococcus aureus* and *Pseudomonas aeruginosa* Infections in Europe–Intensive Care Units, (NCT02413242 ClinicalTrials.gov) (*23*). Between June 2015 and October 2018, the study enrolled a total of 2000 adult subjects within 3 days after ICU admission. To be eligible the patients needed to be on mechanical ventilation at ICU admission and have an expected length of hospital stay ≥48 h. Participants were followed through their ICU stay to assess the development of pneumonia. Data on antibiotic use in the two weeks preceding ICU admission and during the ICU stay were reported. During ICU stay, lower respiratory tract samples were obtained three times in the first week, two times in the three following weeks, on the day of diagnosis of protocol pneumonia and seven days after it. The demographic and clinical baseline characteristics of the 35 subjects included in this analysis are comparable to those of the rest of the study cohort (Supplementary Table 3), except for the APACHE IV score that was lower for the patients included in this analysis.

### Sample collection and isolation

Lower respiratory samples used in this study were collected within the ASPIRE-ICU study (*23*). Untreated respiratory samples were stored at −80 °C until shipment and further analysis at the central lab at the University of Antwerp. The samples were blended (30,000 rpm, probe size 8 mm, steps of 10 s, max 60 s in total), diluted 1:1 v/v with Lysomucil (10% Acetylcysteine solution) (Zambon S.A, Belgium) and incubated for 30 min at 37 °C with 10 s vortexing every 15 min. Thereafter, quantitative culture was performed by inoculating 10-fold dilutions on CHROMID *P. aeruginosa* Agar and blood agar using spiral plater EddyJet (IUL, Spain). Plates were incubated at 37 °C for 24 h and CFU/mL was calculated. Plates without growth were further incubated for 48 h and 72 h. Matrix-Assisted Laser Desorption Ionization-Time of Flight Mass Spectrometry (MALDI-TOF MS) was used to identify 12 *P. aeruginosa* colonies per sample, which were stored at −80 °C until shipment to the University of Oxford and further use.

### Resistance phenotyping

All *P. aeruginosa* isolates were grown from glycerol stocks on Luria-Bertani (LB) Miller Agar plates overnight at 37 °C. Single colonies were then inoculated into LB Miller broth for 18–20 h overnight growth at 37 °C with shaking at 225 rpm. Overnight suspensions were serial diluted to ∼5 × 10^5^ CFU/mL. Resistance phenotyping was carried out as minimum inhibitory concentration (MIC) testing via broth microdilution as defined by EUCAST recommendations (*38, 39*), with the alteration of LB Miller broth for growth media and the use of *P. aeruginosa* PAO1 as a reference strain. Antibiotics were assayed along the following 2-fold dilution series between: ciprofloxacin (0.125 ug/mL - 16 ug/mL), aztreonam (1 ug/mL - 128 ug/mL), ceftazidime (1 ug/mL - 256 ug/mL), meropenem (0.25 ug/mL - 64 ug/mL), piperacillin/tazobactam (2 ug/mL - 256 ug/mL) and gentamicin (0.5 ug/mL - 128 ug/m). Growth inhibition was defined as OD_595_ < 0.200. We calculated a single biologically independent MIC for each of the 441 *P. aeruginosa* isolates on each antibiotic (Supplementary Table 1). When an isolate reached the measurable limit of the MIC assay (i.e. not inhibited at the highest concentration used), the MIC was recorded as double of the upper limit in the raw data file of MIC results (Supplementary Table 1). The number of resistance phenotypes was calculated as the number of MICs per isolate above the following: ciprofloxacin (0.5 ug/mL), aztreonam (16 ug/mL), ceftazidime (8 ug/mL), meropenem (8 ug/mL), piperacillin/tazobactam (16 ug/mL) and gentamicin (8 ug/mL). These points were set from EUCAST guidelines for *Pseudomonas* (v11 breakpoint table, and MIC distributions for *P. aeruginosa* data for gentamicin) (*39*). MDR isolates were defined as isolates with 3 or more resistance phenotypes. For longitudinally sampled patients treated with colistin (patient 6, patient 16, patient 18, patient 32, patient 35), the same protocol was used to determine colistin MIC along the following 2-fold dilution series between: 0.5 ug/mL - 64 ug/mL (Supplementary Table 1), and a colistin resistance phenotype was determined as an MIC above 2 ug/mL (Supplementary Table 2) (*39*).

### Sequencing

All isolates were sequenced in the MiSeq or NextSeq illumina platforms yielding a sequencing coverage of 21X–142X. Raw reads were quality controlled with the ILLUMINACLIP (2:30:10) and SLIDINGWINDOW (4:15) in trimmomatic v. 0.39. Quality controlled reads were assembled for each isolate with SPAdes v. 3.13.1 with default parameters. These assemblies were further polished using pilon v. 1.23 with minimum number of flank bases of 10, gap margin of 100,000, and kmer size of 47. Resulting contigs were annotated based on the P. aeruginosa strain UCBPP-PA14 in prokka v. 1.14.0. Each isolate was typed using the *Pseudomonas aeruginosa* multi-locus sequence typing (MLST) scheme from PubMLST (Last accessed on 11.06.2021). Sixteen isolates were sequenced with the Oxford nanopore MinION platform using the FLO-MIN106 flow-cell and the SQK-LSK109 kit. Raw reads were basecalled using guppy v. 0.0.0+7969d57 and reads were demultiplexed using qcat v. 1.1.0 (https://github.com/nanoporetech/qcat). Resulting sequencing reads were assembled using unicycler v. 0.4.8 (*40*), which used SAMtools v. 1.9 (*41*), pilon v. 1.23 (*42*), and bowtie2 v. 2.3.5.1 (*43*), in hybrid mode with respective illumina reads.

### Variant calling

Paired-ended reads were mapped to the *P. aeruginosa* PAO1 reference genome (NC_002516.2) with Bowtie 2 v2.2.4, and pileup and raw files were obtained by using SAMtools v0.1.16 and PicardTools v1.140, using the Genome Analysis Toolkit (GATK) v3.4-46 for realignment around InDels. From the raw files, SNPs were extracted if they met the following criteria: a quality score (Phred-scaled probability of the samples reads being homozygous reference) of at least 50, a root-mean-square (RMS) mapping quality of at least 25 and a coverage depth of at least 3 reads, excluding all ambiguous variants. MicroInDels were extracted from the totalpileup files when meeting the following criteria: a quality score of at least 500, a RMS mapping quality of at least 25 and support from at least one-fifth of the covering reads (*44*). Filtered files were converted to vcf and SNPs and InDels were annotated with SnpEff v4.2. (*45*). Gene absence was evaluated using SeqMonk (https://www.bioinformatics.babraham.ac.uk/projects/seqmonk/) and OprD structural integrity was investigated within the *de novo* assemblies using an appropriate reference sequence. Finally, all mutations within a set of genes known to be involved in antibiotic resistance were extracted and natural occurring polymorphisms were filtered (*46*). The presence of horizontally acquired antimicrobial resistance determinants was also investigated using the web tool ResFinder (https://cge.cbs.dtu.dk/services/ResFinder/).

To identify mutations and gene gain/loss during the infection, short-length sequencing reads from each isolate were mapped to each of the four long-read de novo assemblies with bwa v. 0.7.17 using the BWA-MEM algorithm. Preliminary SNPs were identified with SAMtools and BCFtools v. 1.9. Low-quality SNPs were filtered out using a two-step SNP calling pipeline, which first identified potential SNPs using the following criteria: 1. Variant Phred quality score of 30 or higher, 2. At least 150 bases away from contig edge or indel, and 3. 20 or more sequencing reads covering the potential SNP position. In the second step, each preliminary SNP was reviewed for evidence of support for the reference or the variant base; at least 80% of reads of Phred quality score of 25 or higher were required to support the final call. An ambiguous call was defined as one with not enough support for the reference or the variant, and, in total, only one non-phylogenetically informative SNP position had ambiguous calls. Indels were identified by the overlap between the HaplotypeCaller of GATK v. 4.1.3.0 and breseq v. 0.34.0. The variable genome was surveyed using GenAPI v. 1.098 based on the prokka annotation of the short-read de novo assemblies. The presence or absence of genes in the potential variable genome was reviewed by mapping the sequencing reads to the respective genes with BWA v.0.7.17.SNPs/indels.

### Growth assays

All isolates were grown from glycerol stocks on LB Miller Agar plates overnight at 37 °C. Single colonies were then inoculated into LB Miller broth for 18–20 h overnight growth at 37 °C with shaking at 225 rpm. Overnight suspensions were serially diluted to an OD_595_ of ∼0.05 and placed within the inner 60 wells of a 96-well plate equipped with a lid. To calculate growth rate, isolates were then grown in LB Miller broth at 37 °C and optical density (OD595nm) measurements were taken at 10-min intervals in a BioTek Synergy 2 microplate reader set to moderate continuous shaking. Growth rate was then calculated as the maximum slope of OD versus time over an interval of ten consecutive readings, and at least three biologically independent replicate cultures were measured for all of the pulmonary isolates to calculate the mean growth rate of each isolate (Supplementary Table 1).

### Statistics

For each patient sample, we calculated the proportion of isolates that were resistant to each antibiotic as described above (*38, 39*). Changes in resistance for each antibiotic were measured as the difference in proportion of resistant isolates between the final and initial sample for longitudinally sampled patients (Supplementary Table 2). To test drivers of antibiotic resistance we used an ANOVA that included main effects of initial proportion of resistant isolates (continuous variable, 1DF), response type (direct or collateral, with the inclusion of colistin for colistin-treated patients; 1DF), pathogen diversity (monoclonal or polyclonal; 1DF), and we nested patient within pathogen diversity (1DF). We also included an interaction term between initial proportion of resistant isolates and pathogen diversity (1DF).

Fitness costs were assessed by comparing the growth rate of co-occuring MDR (i.e. 3 or more resistance phenotypes) and non-MDR (i.e. 0-2 resistance phenotypes) lung isolates from the same patient. We considered all patients with multiple (i.e. >1) MDR and non-MDR isolates in this analysis, giving a total of 179 isolates from 10 patients (Figure 4A). To understand the sources of variation in fitness, we used an ANOVA that included main effects of resistance phenotype (ie MDR or non-MDR; 1DF), pathogen diversity (monoclonal or polyclonal; 1DF) and patient nested within pathogen diversity (8DF). We tested for variation in fitness trade-offs between monoclonal and polyclonal populations by including a resistance phenotype*pathogen diversity interaction in the model.

We tested the association between the number of antibiotic resistance-associated SNPs and the type of infection (polyclonal or monoclonal) using a type 2 two-way ANOVA where the dependent variable is the number of resistance SNPs, and the independent variables are the overall number of SNPs and the type of infection (Figure 3B). We also tested the association of either the overall number of SNPs or the number of regions with variable genetic content and the type of infection. We used two type 2 two-way ANOVAs: for the first one, the dependable variable was the total number of SNPs and the independent variables were the number of isolates and the type of infection; for the second one, the dependable variable was the total number of regions with variable genetic contents and the independent variables were the number of isolates and the type of infection

## Supplementary Information

**Supplementary Table 1: Dataset of 441 *P. aeruginosa* isolates**.

Legend – Metadata of 441 *P. aeruginosa* isolates in study, indicating outputs of genome analysis (including ST), phenotyping (mean growth rate, MICs, resistance phenotyping, MDR classification) and patient information (including monoclonal or polyclonal population classification).

**Supplementary Table 2: Collateral or direct treatment response type to antibiotic treatment**.

Legend – Antibiotic treatment information for monoclonal and polyclonal longitudinally sampled patients, showing antibiotic treatment predicted active for *Pseudomonas* (EUCAST guidelines). Collateral or direct treatment response type was determined from MIC screening against the following antibiotics (/antibiotic classes): meropenem (used as direct response type for patients treated with carbapenems), ciprofloxacin (used as direct response type for patients treated with fluroquinolones), ceftazidime (used as direct response type for patients treated with cephalosporins), gentamicin (used as direct response type for patients treated with aminoglycosides), aztreonam, piperacillin/tazobactam and colistin. Initial_medianMIC_; Median MIC value for isolates from the first sampling point Final_medianMIC_; Median MIC value for isolates from the final sampling point Initial_R_: Proportion of resistance phenotypes in isolates from the first sampling point Final_R_: Proportion of resistance phenotypes in isolates from the final sampling point

**Supplementary Table 3: Comparison of characteristics of study cohort and total ASPIRE-ICU set**.

**Supplementary Figure 1: Within-strain genetic diversity of monoclonal and polyclonal populations**.

Legend - Frequency of (A) all variants, SNPs and indels, and (B) regions of variable genetic content in monoclonal and polyclonal populations. There was no significant difference ([A] F_1,67_=3.742, P=0.063, [B] F_1,67_=0.183, P=0.672) in the distribution of these variants between monoclonal and polyclonal infections.

## Acknowledgements

This research was supported by Wellcome Trust Grant (106918/Z/15/Z) and the Innovative Medicines Initiative Joint Undertaking under COMBACTE-MAGNET (Combatting Bacterial Resistance in Europe-Molecules against Gram-negative Infections, grant agreement no. 115737) and COMBACTE-NET (Combatting Bacterial Resistance in Europe-Networks, grant agreement no. 115523), resources of which are composed of financial contribution from the European Union’s Seventh Framework Program (FP7/2007-2013) and EFPIA companies’ in kind contribution. We thank the Oxford Genomics Center (funded by Wellcome Trust Grant 203141/Z/16/Z) for the generation and initial processing of Illumina sequence data.

## Notes

### Competing Interest Statement

The authors have declared no competing interest.

